# Atomistic Mechanisms of Calcium Permeation Modulated by Q/R Editing and Selectivity Filter Mutations in GluA2 AMPA Receptors

**DOI:** 10.1101/2024.12.03.626525

**Authors:** Florian Heiser, Johann Biedermann, Ece Kuru, Andrew J. R. Plested, Han Sun

## Abstract

GluA2 is a key subunit of AMPA receptor ion channels that is abundantly expressed in the vertebrate brain. Post-transcriptional Q/R editing of GluA2 renders AMPARs nearly impermeable to calcium ions, which is crucial for their normal function. Although previous studies have characterized conductivity and selectivity differences between edited and unedited GluA2 variants and heteromeric receptors incorporating GluA2, the atomistic mechanism remains largely unknown. In this study, we investigate ion permeation in the context of multiple Ca^2+^ binding sites along the pore predicted from MD simulations, considering both mutations and co-permeating monovalent ions. Patch clamp electrophysiology recordings confirmed a binding site at the intracellular mouth of the selectivity filter that confers selectivity for calcium over monovalent ions. A patient mutation at the same site has been previously shown to cause neurodevelopmental abnormalities. Furthermore, MD simulations of GluA2 with different arginine copy number at the Q/R site show that Ca^2+^ conduction is blocked in the presence of two arginines, whereas K^+^ is only blocked by four arginines, in explaining the results from decades of electrophysiological work. Finally, MD simulations revealed that Ca^2+^ reduces K^+^ conduction by preferentially occupying the intracellular SF binding site, whereas Na^+^ does not. This result is consistent with electrophysiological results from the D590 mutants, and suggests that divalent binding in the selectivity filter is a major determinant of AMPAR conductance.

## Introduction

Glutamate receptor ion channels are abundant in the vertebrate central nervous system, and the *α*-amino-3-hydroxy-5-methyl-4-isoxazolepropionic acid receptor (AMPAR) subtype plays an essential role in mediating fast excitatory neurotransmission. At the synaptic cleft, when glutamate binds to the ligand binding domain, the transmembrane ion channel domain of the AMPAR opens, allowing the passive flux of K^+^, Na^+^ and Ca^2+^ ions across the postsynaptic membrane. This event results in a brief depolarization of the postsynaptic neuron. In contrast to sodium and potassium, calcium is a second messenger that has wide ranging roles in cellular excitability, neuronal development and the plasticity of neural circuits. Therefore, the regulation of calcium permeability of AMPA receptors is vital, and is determined by the presence or absence of the GluA2 subunit. In all AMPA receptor subunits (GluA1-4) a glutamine residue is encoded in genomic DNA at the top of the selectivity filter (SF, residue 586, Figure 1). But in GluA2 this codon undergoes post-transcriptional RNA editing (1) and is translated as arginine (the so-called Q/R editing site) in *>* 95 % of transcripts in the adult brain (2). Reduced Q/R editing is deleterious for brain function (3, 4). Combination of the GluA2(R) subunit with other GluA subunits (which are not edited and therefore provide Q residues) renders the AMPAR relatively impermeable to calcium ions (calcium impermeable; CI-AMPARs, (5, 6)). A receptor composed entirely of GluA2(R) has a very low conductance for monovalent cations (7). However, a long-standing unresolved question is, what is the relation between conductance, calcium permeability and the copy number of arginines in the pore (8, 9). How many arginines are needed to prevent calcium flux, and what is the molecular mechanism?

**Fig. 1.**
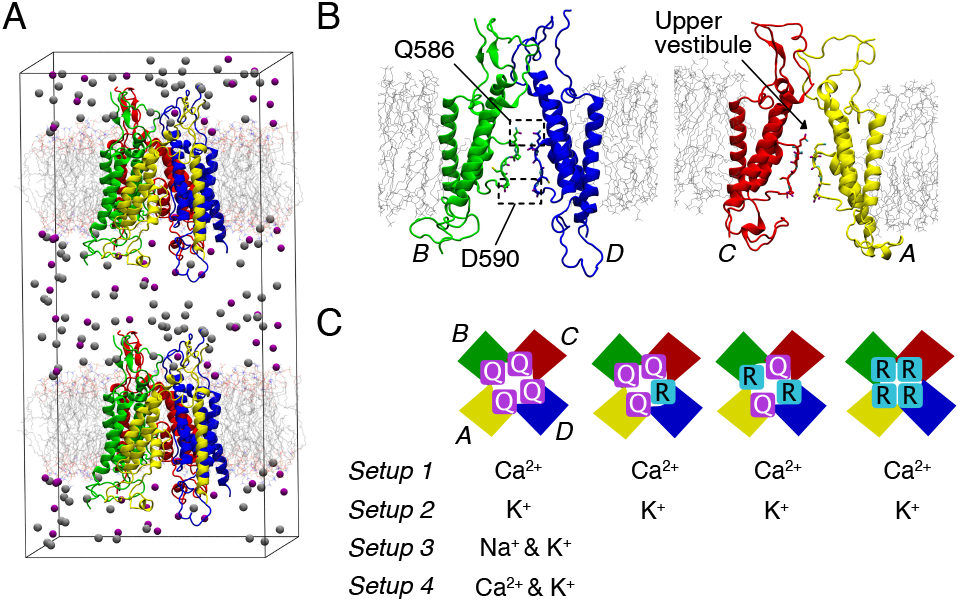
Computational setup. (A) MD-based computational setup for simulating ion co-permeation in the open-state of GluA2 (PDB ID: 5WEO). Only the Transmembrane Domain (TMD) and linkers to the Ligand Binding Domain (LBD) were included in the simulations, following previously developed strategies (27, 28). (B) A/C (left) and B/D (right) subunits with the Q586 at the Q/R editing site and D590 on the intracellular side of the SF labeled. Upper vestibule between the Q/R site and the upper gate is indicated with an arrow. (C) Four different computational setups with varying Q/R arrangements and ion/ion mixture types.

Recent advancements in cryogenic electron microscopy (cryo-EM) have yielded invaluable maps of the structural changes of various AMPAR isoforms, providing a deeper understanding of their gating process in closed, partially open, and fully open states (10–20). These studies led to a proposal that the Q/R-editing site in the selectivity filter (SF) acts as gate, perhaps providing an explanation for how auxiliary subunits might enhance channel conductance (21). Other work also suggests coupling between the selectivity filter and gating, but related to desensitized states (22, 23). Notably, recent high-resolution cryo-EM structures of AMPARs revealed a Ca^2+^ binding site towards the extracellular side, which is selective for Ca^2+^ over Na^+^ and was proposed to be functionally important for Ca^2+^ permeation (24). Moreover, a cryo-EM study of native AMPA receptor complexes by Zhao and Gouaux et al. unveiled preferred combinations of subunit arrangements, with GluA2 being the most abundant single subunit (25), being most often found in two of the four possible sites, diagonally opposed. Despite the findings from these studies, one disadvantage of purely structural insights is that they are (usually) performed in the absence of transmembrane voltage. Thus, a gap remains in our atomistic understanding of the relationship between Ca^2+^ permeation in non-equilibrium conditions, and GluA2 incorporation. With improvements in force field models and advancements in computational power, molecular dynamics (MD) simulations have evolved into a powerful theoretical method for investigating ion permeation, ligand modulation and the gating dynamics of ion channels (26). Following the release of the first open cryo-EM structures of GluA2, we conducted extensive MD simulations of GluA2 involving various cation species. These simulations suggested that the permissive structure of the SF can accommodate different monovalent cations at a single site, each retaining their distinct hydration state during ion permeation (27). More recently, we performed an integrated study to investigate the Ca^2+^ permeation mechanism in a Ca^2+^-permeable AMPAR (CP-AMPAR) variant of GluA2 (28). Compared to monovalent cation permeation, Ca^2+^ occupies multiple binding sites along the ion permeation pathway. Notably, MD simulations predicted a stable Ca^2+^ binding site on the intracellular side of the SF (CDI region), which could be introduced to the NaK channel by mutation as confirmed by crystallography. The physical chemistry of this site was validated by ab initio quantum mechanics/molecular mechanics (QM/MM) calculations.

In the present study, we confirmed the essential role of the Ca^2+^ binding site on the intracellular side of the SF for Ca^2+^ permeability, as predicted by MD simulations. Mutation of the aspartic acid 590 in the filter leads to a positive shift in the reversal potential in patch clamp recordings with Ca^2+^ relative to Na^+^. Furthermore, we conducted a large number of all-atom MD simulations to model Ca^2+^ and K^+^ permeation in GluA2 while varying the Q/R editing levels and employing either single or mixed ion types (Figure 1C). Our results demonstrated distinct conductance properties, binding sites, and pathways for K^+^ and Ca^2+^, highlighting the ability of the intracellular Ca^2+^ binding site in the SF for blocking K^+^ conduction. This study provides valuable atomistic insights into cation selectivity, the relationship between Q/R editing levels and cation conductance, and the mechanism by which calcium blocks co-permeant monovalent cations.

## Results

### Binding of Ca^2+^ in the gate region

Recently high-resolution cryo-EM structures of GluA2 were reported showing a Ca^2+^ binding site, *site-G*, proposed to be critical in regulating ion transport (24). The authors suggested that *site-G* was distinct from binding sites identified in our previous MD simulations (28). Consequently, we systematically compared the location of *site-G* with the ion occupancy and trajectories from these previous simulations in a pure calcium condition. As shown in Figure 2 and Movie S1, *site-G* was well covered by our previous MD simulations. However, since *site-G* and binding site 4 are very close to each other, we had not differentiated them in our previous classification. In Movie S1 it is evident that Ca^2+^ frequently exchanges between *site-G* and binding site 4, indicating that binding is not particularly stable at either site under transmembrane voltage. In simulations, *site-G* and site 4 form a diffuse, bottle-shaped region of ion occupancy, albeit with lower occupancy than the QQ site and the upper vestibule (Figure 2).

**Fig. 2.**
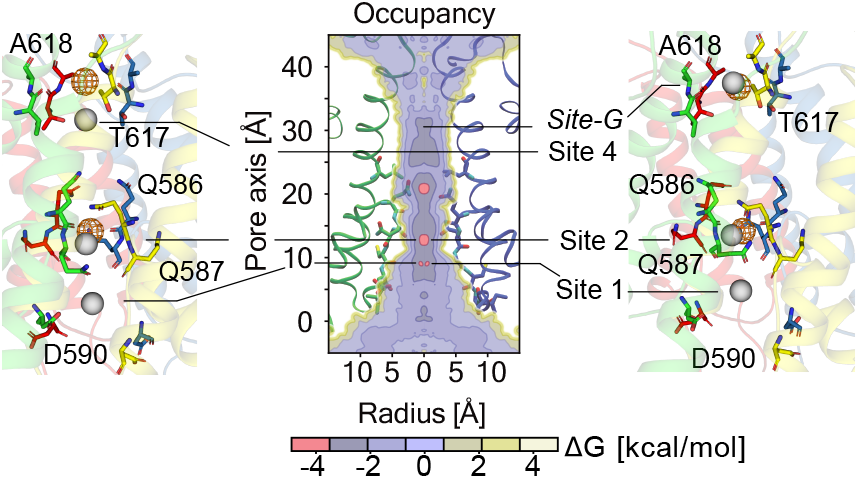
Comparison of calcium ion binding sites determined by MD simulations and cryo-EM. Two selected snapshots (left and right panels) from our previous MD simulations (28) of GluA2(Q) showing Ca^2+^ ions at their binding sites (grey spheres) during ion permeation, overlaid onto the Ca^2+^ binding sites (orange mesh) determined from a high-resolution cryo-EM structure of AMPAR (PDB ID: 8FQ1) (24). AMPAR subunits are displayed as ribbons and selected residues in liquorice. Ion binding sites identified in our MD simulations, along with the experimentally-resolved *site-G*(24), are labeled on the two-dimensional plot of Ca^2+^ occupancy (middle, modified from ref. (28)). In unedited receptors, Ca^2+^ also shows a strong occupancy at a site just above the Q/R site (not labelled here). The ion occupancy is color-coded in terms of the free energy difference to the bulk solution outside the channel.

To reconcile these differences, it is important to consider three major distinctions in the way ion binding sites have been derived from MD simulations and experimental characterization by cryo-EM structures: (i) MD simulations provide a dynamic picture of ion permeation, where ion occupancy reflects a statistical superimposition sampling all major and minor binding sites during the permeation process. This contrasts with cryo-EM structures, which might select very stable ion binding sites along the ion permeation pathway under equilibrium; (ii) The simulations were performed under transmembrane voltages, mimicking membrane potentials under non-equilibrium conditions. In contrast, in standard cryo-EM experiments, structures are resolved under zero membrane potential; (iii) Our simulations were performed in the absence of flanking Stargazin molecules, which were present for structure determination and could alter the ion binding sites, because they are known to alter pore structure from their effects on polyamine block and conductance (29, 30).

### Mutation at D590 reduces Ca^2+^ permeability and block

Notwithstanding the differences in interpretation around *site-G*, a more substantial difference between our previous MD simulations of Ca^2+^ permeation (28) and the high-resolution cryo-EM structures analyzing Ca^2+^ binding in GluA2 (24) is found within the SF (CDI region, site 1). This binding site shows substantial occupancy for calcium ions in our simulations. In the cryo-EM analysis of Nakagawa et al., this site is marked (as Site-C589) but neither further analyzed nor tested, and appears secondary, with perhaps lower occupancy. Notably, cryo-EM data suggests the selectivity filter is quite dehydrated and shows little Ca^2+^ occupancy even at the QQ (site 2). In our simulations, there is substantial accumulation of Ca^2+^ and water in the region “below” the SF, which we also previously observed in simulations and crystal structures of the CDI mutant of NaK (referred to as site 0 in ref. (28). A previous clinical study identified a mutation at D590 as being responsible for a neurodevelopmental disorder in one patient (numbered as D611N in ref. (31)). To further evaluate this putative Ca^2+^ binding site, we mutated D590 to either asparagine or serine and performed patch clamp electrophysiology experiments in sodium-and calcium-rich buffers.

Both mutants expressed well and gave fast glutamate-activated responses that exhibited similar desensitization to wild-type GluA2 (Figure 3). Upon switching to a high Ca^2+^ solution, the wild-type current voltage relation remained approximately linear but shifted on average 25 mV to a more positive potential, indicating a high relative permeability for Ca^2+^. Without correcting for ion activity (see Methods), the ratio PCa^2+^ : PNa^+^ was 4. Both mutants almost abolished this shift, reducing it to 6 mV (D590N) and 7 mV (D590S), indicating nearly equal permeability of the two ion species (PCa^2+^ : PNa^+^ was 1.15 for D590N and 1.2 for D590S). We note that these ratios are difficult to compare with previous work because of inconsistencies with the calculations that have been used. As a guide, the absence of any shift would in fact correspond to a PCa^2+^ : PNa^+^ of 0.83, that is, calcium being less permeant than sodium, because it has a higher valence.

**Fig. 3.**
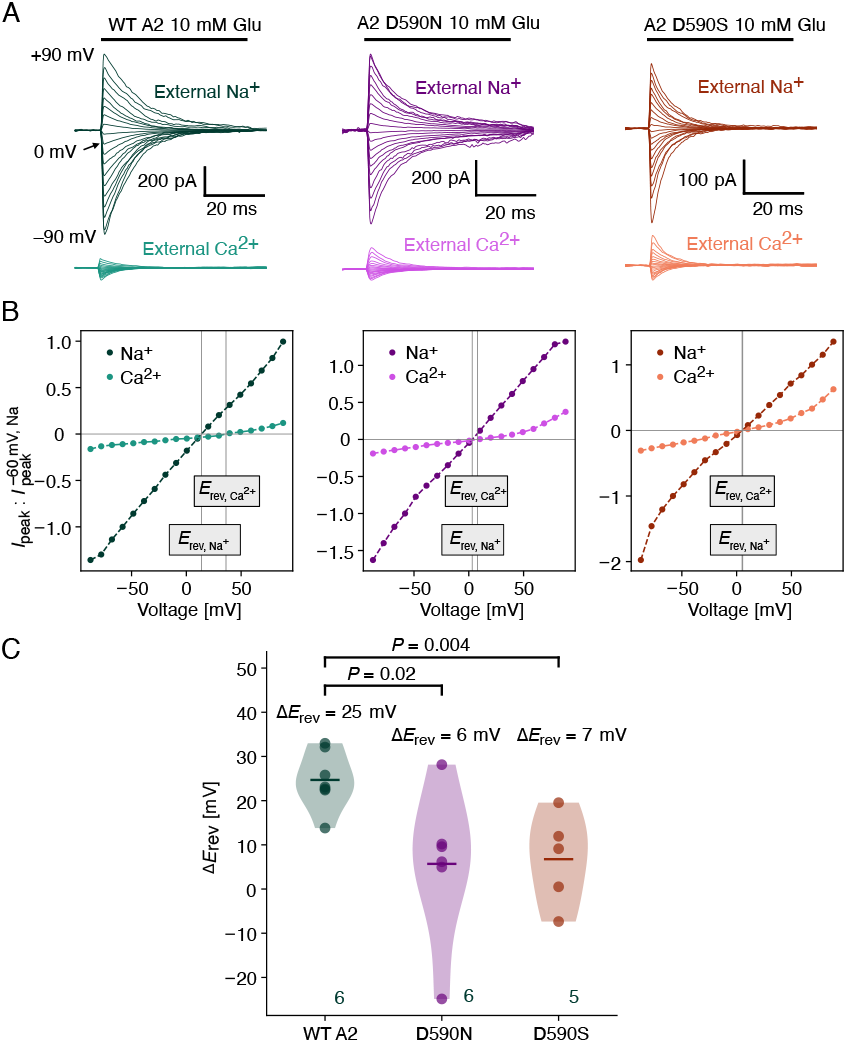
Mutations at Site 1 (CDI region) render Ca^2+^ and Na^+^ equally permeant. (A) Responses to fast jumps into 10 mM glutamate (black bar) over a range of command potentials between –90 mV and +90 mV for wild-type GluA2 and two mutants at site 1, D590N and D590S are plotted. The upper row shows the currents in external Na+ (150 mM) and the lower row shows the smaller responses from the same patch in external Ca^2+^ (100 mM). (B) Current voltage plots show the reversal potentials (E_rev_, grey lines) in each condition for the examples in panel A. (C) The shift in E_rev_ between the two conditions (Na^+^ and Ca^2+^). The number of patches in each group is indicated. The Alexander-Govern test of the equality of the three means gave a P-value of 0.008.

Although the disruption of site 1 (at the CDI region) abolished selective permeability for calcium ions, confirming it to be a major site for Ca^2+^ binding in the channel, the overall conductance in high Ca^2+^ remained similarly low to wildtype for both mutants. At this high concentration of external calcium ions, there is no voltage dependence of the reduced conductance in wild-type GluA2. But both mutants, and in particular the D590S mutant, showed a relief of Ca^2+^ block at potentials highly positive of the Ca^2+^ reversal potential. In this condition, Na^+^ efflux is competing with Ca^2+^ influx, and one interpretation of this observation is that at high Ca^2+^, a lower affinity, voltage-dependent block (which could be due to *site-G* and/or other sites) is revealed by mutation, which ablates the highest-affinity site.

These results suggest that the CDI site in the SF (site 1) is a relatively high-affinity site, perhaps outside the electric field. When it is disrupted, block at high Ca^2+^ becomes voltage-dependent. In narrow regions, electrostatic interactions between permeant ions over space can develop and in principle couple remote sites to the electric field (32). As will be shown in the following, such a coupling between permeating monovalent and divalent ions might be pronounced in the AMPAR pore.

### Cation conductances are anti-correlated with the level of Q/R editing

Our previous simulations have suggested a stable cation binding site at the Q/R editing site in simulations of either monovalent ions or Ca^2+^ (27, 28), while functional electrophysiology has shown that Q/R editing alters the Ca^2+^ permeability of AMPARs (5, 6). Therefore, here we simulated the permeation of K^+^ and Ca^2+^ ions, respectively, at different Q/R editing levels of GluA2 using our previously developed MD-based approach ((27, 28), Fig. 1A). We used the unedited form of the open-state GluA2 (PDB ID: 5WEO) as the structural template and systematically substituted glutamine with arginine in three different configurations (QRQQ, QRQR, and RRRR, (Figure 1C)) in the simulations, and compared these to our previously published simulations on the QQQQ configuration (27, 28). Note that although that all glutamate receptors are overall two-fold symmetric, the transmembrane domain of GluA2 is nearly four-fold symmetric. However, for consistency with heteromeric structures where GluA2 (R) occupies the B and D positions (25), we added arginine residues to these positions in the QRQQ and QRQR configurations.

The results, summarized in Figure 4A, indicated that introducing positively-charged arginine in one subunit (QRQQ) reduces the simulated conductance of K^+^ and Ca^2+^ by 30-50 %. Introducing two arginines completely abolished Ca^2+^ conductance in the simulations, while 20 % of K^+^ conductance remained compared to the unedited form. This matches well expectations from heteromeric AMPARs incorporating GluA2(R) (5, 6). In the fully edited form of GluA2(RRRR), only a very limited amount of K^+^ permeation was observed. This result is similar to experimental single-channel recording of homomeric GuA2(R) channel, revealing a dramatically reduced unitary conductance for monovalent cations (7). Note, similar to our previous simulations (27, 28), we mostly observed the outward cation permeation. Therefore, we only compared the outward permeation across different simulation setups.

**Fig. 4.**
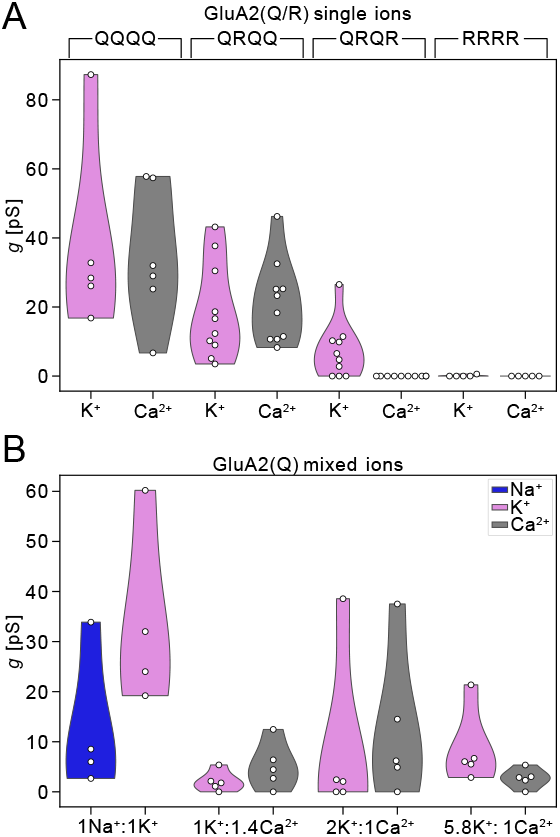
Comparison of simulated conductances for single and mixed cation types. (A) Simulated K^+^ (magenta) and Ca^2+^ (grey) conductances derived from the simulations in QQQQ, QRQQ, QRQR, and RRRR setups. The simulations were performed with 180-212 mM KCl and 90 mM CaCl2, respectively. Five runs of 250 ns simulations were performed for each setup, at 303 K and 518 *±* 76 mV using the C_HARMM_36 force field. Only the outward conductance is shown. The data from the QQQQ setups (Na^+^ and Ca^2+^, each alone) are taken from our previous studies (27, 28). (B) Simulated K^+^ (magenta), Na^+^ (blue), and Ca^2+^ (grey) conductances derived from the simulations with the unedited AMPAR (QQQQ) with mixed cations in each simulation setup. Five runs of 250 ns simulations were performed for each setup, at 303 K and 485 *±* 120 mV using the C_HARMM_36 force field.

Analysis of the two-dimensional ion occupancy (Figures 5 and 6) suggested that increasing the number of arginines at the Q/R site gradually decreases cation occupancy in the upper vestibule, that is, the region between the Q/R editing site and the upper gate (Figure 1B). This reduction is expected due to the electrostatic effect of the positively charged arginines in the narrow pore. Even a single replacement of glutamine with arginine (QRQQ setup) led the occupancy of the external binding site slightly below the upper gate (site 4) to vanish for K^+^ and substantially reduce in occupancy for Ca^2+^. In the QRQR simulations in pure Ca^2+^, where no ion conduction was observed, the entire vestibule region was devoid of Ca^2+^ ions, providing a plausible explanation for the absence of conductance. In the fully edited form of GluA2(RRRR), Ca^2+^ ions could only reach the intracellular binding site in the SF, unable to approach the positively charged ring of guanidinium groups formed in the vicinity of the extracellular side of the filter.

**Fig. 5.**
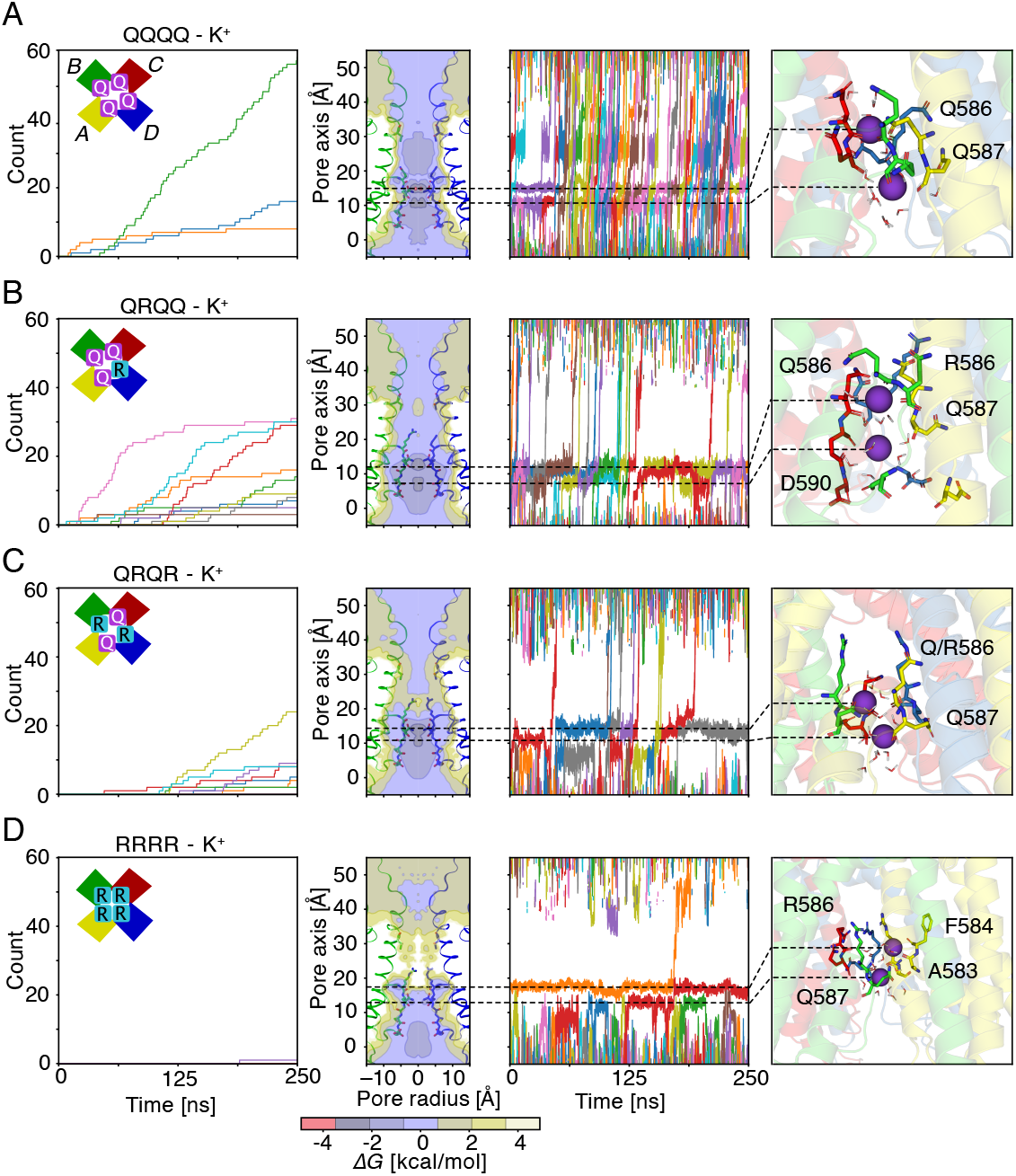
K^+^ permeation in edited (QRQQ, QRQR, RRRR) and unedited (QQQQ) forms of GluA2. Comparison of the cumulative number of outward K^+^ permeations during five runs of 250 ns simulations with 180 mM KCl (column 1), two-dimensional K^+^ ion occupancy along the ion permeation pathway (column 2), representative traces of K^+^ passing through the SF of the GluA2 channel pore (column 3), and a representative snapshot of K^+^ binding (column 4) in (A) QQQQ, (B) QRQQ, (C) QRQR, and (D) RRRR setups. Note, that 10 simulation replicates were performed for QRQQ and QRQR, while 5 simulation replicates have been carried out for all other setups. Also, the RRRR setup was simulated for 500 ns while only 250 ns are shown in the ion track plot. The data from the QQQQ setup is taken from our previous study (27).

**Fig. 6.**
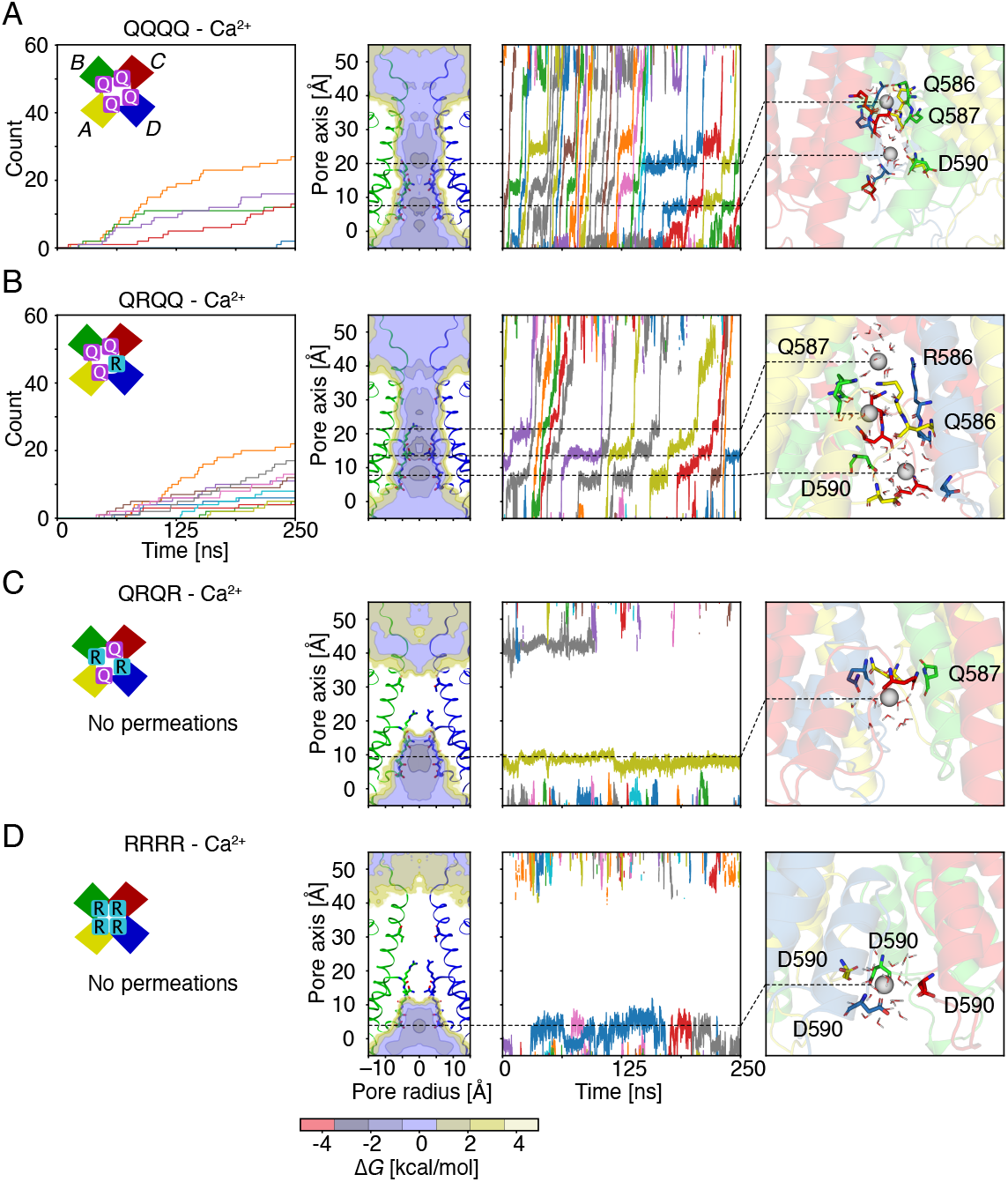
Ca^2+^ permeation in edited (QRQQ, QRQR, RRRR) and unedited (QQQQ) forms of GluA2. Comparison of the cumulative number of outward Ca^2+^ permeations during five runs of 250 ns simulations with 90 mM CaCl2 (column 1), two-dimensional Ca^2+^ ion occupancy along the ion permeation pathway (column 2), representative traces of Ca^2+^ passing through the SF of the GluA2 channel pore (column 3), and a representative snapshot of Ca^2+^ binding (column 4) in (A) QQQQ, (B) QRQQ, (C) QRQR, and (D) RRRR setups. Note, that 10 simulation replicates were performed for QRQQ and QRQR, while 5 simulation replicates have been carried out for all other setups. The data from the QQQQ setup is taken from our previous studiy (28), but without splitting sections according to high and low conductance.

In contrast, increasing the number of arginines and thus adding positive charges above the selectivity filter had a smaller effect on monovalent K^+^ ions, and K^+^ conductance was retained in the QRQR simulations. Despite weak occupancy of site 4, some K^+^ ions were still able to pass through the channel. Strikingly, in the edited forms of QRQR and RRRR (where the conductance was very low), in the majority of cases, K^+^ ions used an unexpected alternative route to pass the Q/R site. Instead of passing through the canonical pathway, along the central symmetrical axis, the K^+^ ions used a side pocket “behind” the SF, coordinated by A583 and F584 (Figure 5D). This path is witnessed by the 2D ion occupancy, where the ion density extends to the lateral region “behind” the SF (see also movies S2 and S3 for an ion transiting this region). In contrast, this side route was not observed for Ca^2+^, possibly because its double positive charge prevented it from being accommodated. Instead, Ca^2+^ primarily stops at the lower binding sites in the filter, coordinated by Q587, C589 and D590 (Figure 6C & D). This difference between K^+^ and Ca^2+^ in utilizing the side route may further explain why monovalent ions can permeate through the channel in the double R-edited and fully edited forms, whereas Ca^2+^ cannot.

Based on structural analyses, it was proposed that a second, lower gate in the channel is formed by sidechains at the Q/R site, which protrude towards the centre of the ion channel pore in the otherwise open structure (14). We calculated the minimum distance between the side-chain of R586 and the centre of the ion channel pore in the QRQQ, QRQR, and RRRR simulation setups (Figure S1). While the side-chain of R586 was highly dynamic during simulations, we did not observe any correlation between conduction and the postion of R586. Therefore, our simulations did not support the hypothesis of a simple gate formed by the Q(R)586 residue. However, as our data indicate, in edited channels, even when the canonical axial path is unavailable, monovalent cations can take another route and so there is no simple relation between the Arginine positions and conductance.

### K^+^ and Na^+^ co-permeate quasi-independently through the GluA2(Q)

We further performed MD simulations with mixed monovalent cations, K^+^ and Na^+^. The total K^+^ and Na^+^ concentration was 212 mM, similar to our previous simulations with individual K^+^ and Na^+^ ions (27). These simulations yielded concurrent conductances of 34 *±* 16 pS for K^+^ and 13 *±* 12 pS for Na^+^ (Figure 2C), which are similar to the conductances derived from simulations with a single monovalent cation type (K^+^ : 38 *±* 25 pS and Na^+^ : 14 *±* 7 pS) (27). Analysis of 2D ion occupancy and representative ion track plots of K^+^ and Na^+^ passing through the filter suggested that these two monovalent ion types occupy similar ion binding sites within the filter during their permeation (Figure S1). Therefore K^+^ and Na^+^ co-permeate GluA2(Q) following a common monovalent ion mechanism, with efficient knock-on between ions, irrespective of which species is involved. Importantly, K^+^ and Na^+^ neither impede each other, nor do they promote new “coupled” ion occupancy features, where groups of ions (e.g. pairs of K^+^ ions) interact preferentially, suggesting a form of independence between monovalent ions. As shown below, this is in distinct contrast with K^+^ and Ca^2+^.

### Ca^2+^ blocks co-permeating K^+^ conduction by occupying the intracellular SF binding site

Next, we performed ion permeation simulations at three different Ca^2+^ and K^+^ ratios. The total cation concentration was 120 - 180 mM. Notably, even with a large excess of K^+^ over Ca^2+^ at a ratio of 5.8 : 1, we observed a 5-fold reduction in the computed K^+^ conductance to 8 *±* 7 pS (compared to our previous K^+^-only simulations, 38 *±* 25 pS, at similar total cation concentration). In this setup, a small but consistent Ca^2+^ conductance of 2 *±* 2 pS was discernible. When the Ca^2+^ proportion was increased to half that of K^+^ (K^+^2 : Ca^2+^ 1), conductances for both ions rose, but only weakly. When the proportion of Ca^2+^ was further increased, to a condition where Ca^2+^ was in excess, both ion conductances remained small. In none of the three conditions were the conductances measurably different from one another, and conductances computed over each sub-microsecond simulation duration showed a large variance (figure 7C).

**Fig. 7.**
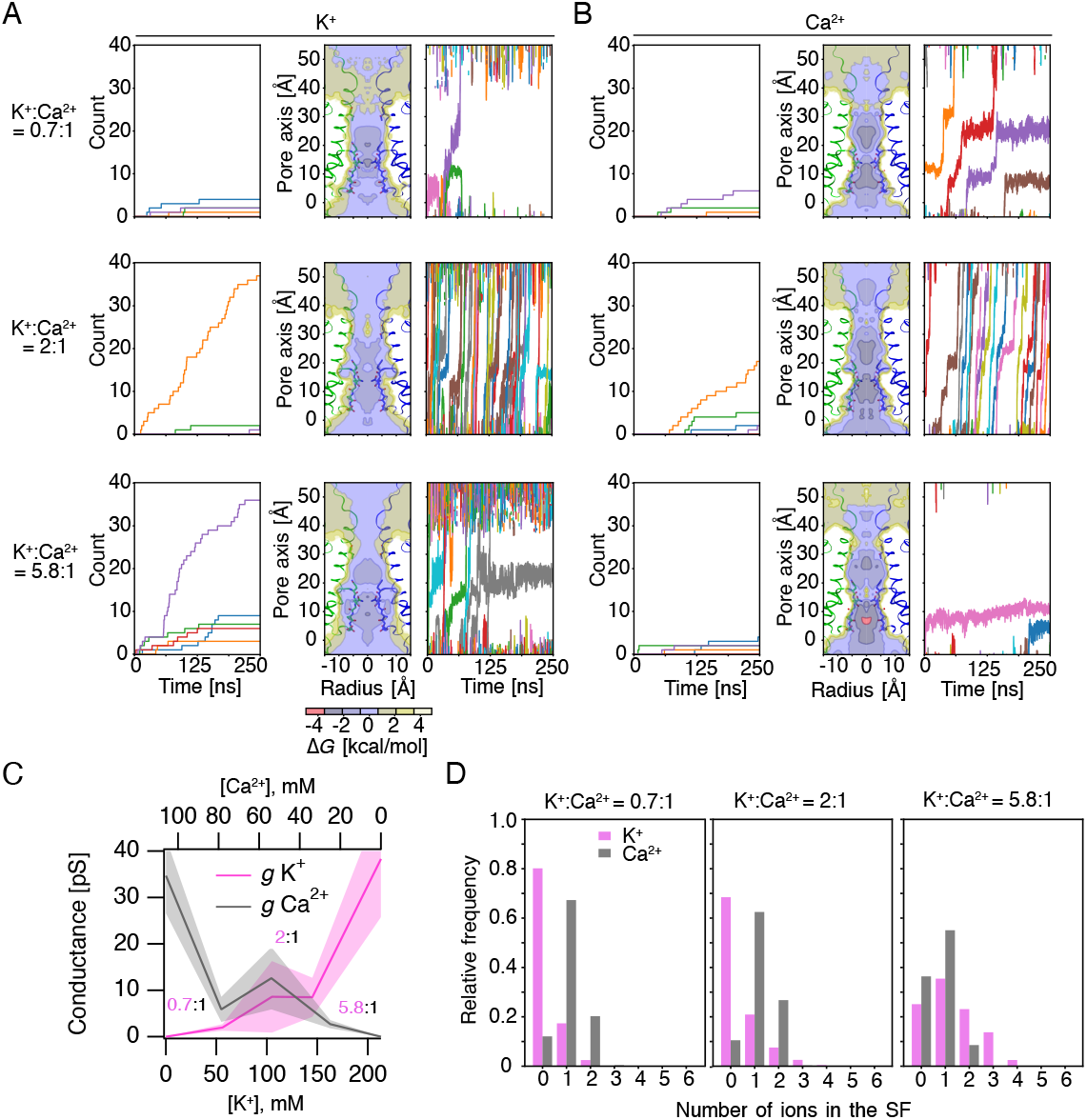
Mixed cation permeation in unedited form (QQQQ) of GluA2. (A) Comparison of the cumulative number of outward K^+^ permeations (column 1) two-dimensional K^+^ (column 2) occupancy along the pore, and representative traces of K^+^ (column 3) passing through the SF of the GluA2 channel pore in (upper row) 0.7 K^+^ : 1 Ca^2+^, (middle row) 2 K^+^ : 1 Ca^2+^, and (lower row) 5.8 K^+^ : 1 Ca^2+^ simulations. Ion occupancy is plotted on a logarithmic scale as concentration, and linearly as free energy. The total K^+^/Ca^2+^ concentration is 120-180 mM. (B) The results from the same representative simulations are shown for the co-permeating calcium ions. (C) Comparisons of the mean computed conductances (*g*) for K^+^ and Ca^2+^ across the three mixed conditions. Five runs of 250 ns simulations are shown for each setup with C_HARMM_36 force field. The 5.8:1 setup was simulated for 500 ns but only the first 250 ns are shown in the ion track plot. (D) Population of number of K^+^ and Ca^2+^ in the SF during the simulations with 1 K^+^ : 1.4 Ca^2+^ (left), 2 K^+^ : 1 Ca^2+^ (middle), and 5.8 K^+^ : 1 Ca^2+^ (right).

Analysis of 2D ion occupancy and K^+^ and Ca^2+^ ion traces passing through the filter suggested that at low Ca^2+^ proportion, Ca^2+^ ions preferentially bind to the intracellular binding site (site 1) in the SF, substantially reducing K^+^ permeation (Figure 7A&B). This effect is not so different to what is observed for pure calcium permeation (28), where for at least a fraction of the time, Ca^2+^ blocks the channel rather than permeating. These observations provide a plausible mechanism for our electrophysiology measurements, which showed that mutation of D590 leads to a strong reduction in Ca^2+^ selectivity over Na^+^. In the Ca^2+^- and K^+^-only simulations, K^+^ and Ca^2+^ permeate through the filter via a loosely-coupled knock-on mechanism. The mixed ion simulations revealed that the knock-on conduction, whereby one ion “pushes” another one through the narrow region of the channel (33), is rather inefficient between monovalent and divalent cations. Furthermore, the small size and double charge of Ca^2+^ ions allow them to occupy the negatively charged environment of SF (CDI region) more easily than K^+^. This is highly evident in the simulations with a 0.7 K^+^ : 1 Ca^2+^ ratio, showing a strong reduction of K^+^ occupancy in the SF compared to the simulations with higher K^+^ to Ca^2+^ ratio.

To investigate further, we calculated the relative frequency of a given number of K^+^ and Ca^2+^ ions being found in the SF during each simulation setup (Figure 7D). In the 5.8 K^+^ : 1 Ca^2+^ simulation setup, K^+^ and Ca^2+^ were almost equally populous in the SF. However, in the simulations at the two other ratios with increased Ca^2+^ concentration, configurations without K^+^ in the filter dominated. This result again suggests that Ca^2+^ outcompetes K^+^ for occupation of the filter. Interestingly, the maximum number of charges the SF can accommodate seems to be four. In the simulations of 5.8 K^+^ : 1 Ca^2+^, a maximum of four K^+^ ions could occupy the filter, while in the simulations with excess Ca^2+^, the maximum number of Ca^2+^ accommodated in the filter is two.

## Discussion

The regulation of calcium permeation in AMPARs is a crucial process in neurons. In most AMPARs, the GluA2 subunit is edited to arginine at the Q/R site, which reduces calcium influx. On the other hand, calcium-permeable AMPA receptors (CP-AMPARs) lack GluA2, and calcium permeation in the presence of four glutamine residues at the Q/R site permits calcium signaling pathways that modulate synaptic strength (34). Transient insertion of GluA1 (or CP-AMPARs in general) may also underlie plasticity (35, 36). In interneurons, CP-AMPARs lacking GluA2 are the predominant form (37). The degree to which CP-AMPARs contribute alongside GluA2-containing CI-AMPARs in principal neurons remains unclear (38–40). Aside from these considerations, the atomistic mechanisms of calcium permeation or rejection in AMPARs remains under debate. In this study, we simulated the permeation of Ca^2+^, K^+^, and a mixture of different monovalent and divalent cation types through the GluA2 channel pore with various Q/R configurations. Mixtures represent the physiological condition, where high external sodium and high internal potassium compete with calcium influx. We propose that, compared to monovalent cations, Ca^2+^ ions are more sensitive to mutations and residue editing along the ion permeation pathway, as Ca^2+^ ions exhibit multiple binding sites within the SF, as well as between the SF and the M3 gate in AMPAR.

A previous structural investigation and functional experiments on a neighbouring mutant residue focussed on an extracellular binding site (*site-G*) at the channel gate region and the Q/R site for Ca^2+^ transport through the AMPAR pore (24, 41). Specifically, it was proposed that *site-G* is the only Ca^2+^ blockage site for monovalent cation conduction. This idea contrasts with findings from our previous MD simulations, and what we present here. In the present study, we experimentally validated the functional role of an additional stable binding site, which was predicted by MD simulations. This site is located on the intracellular side of the SF (CDI region). Notably, the negatively charged, highly conserved aspartic acid residue among the different GluA variants contributes to the binding of Ca^2+^. In kainate and NMDA subtype glutamate receptors (GluK and GluN), a negatively charged glutamic acid replaces aspartic acid, but in principle these could similarly form a divalent cation binding site. Moreover, we showed here that mutations of aspartic acid to asparagine or serine in GluA2 led to an abolition of the positive shift in the reversal potential and concomitant reduction of calcium selectivity, and a reduction in block in biionic conditions (Figure 3). This position (often called +4 because it is 4 residues after the Q/R site) was shown to be important for polyamine block and divalent ion binding (42, 43), as well as the action of Stargazin on conductance (30), but received little attention in a structural investigation of the GluA2 pore in the presence of Stargazin (24). Another explanation is that, for a structural study at 0 mV, the ion binding sites have a strikingly different occupancy. For example, perhaps site-G is preferentially occupied in the absence of an external electric field. However, our own work on NaK shows that there is no fundamental reason not to detect either monovalent or divalent ions in the CDI region of the selectivity filter (28, 44) at 0 mV. Finally, Stargazin could influence the divalent ion occupancy of the filter, although Stargazin itself does not affect calcium permeability much (29). A clinical study suggested that a patient with a neurodevelopmental disorder carried the same mutation of D590N (31). Therefore, we suggest an atomistic mechanism for this disease-related mutation, related to the alteration of calcium permeability and ion selectivity in neurons. It will be important in future work to weigh these effects against a possible change of polyamine block (42) in the DN mutant. Further, we propose that, in the presence of a physiological mixture of Ca^2+^ and monovalent ions, permeation in the AMPAR channel is a complex process involving multiple transient cation binding sites along the pore. This differs from the previous view that only one or two binding sites are critical for cation transport in AMPARs.

When simulating GluA2 with different levels of Q/R editing, we observed a reduction of K^+^ and Ca^2+^ conductance when only one glutamine was replaced by arginine, while replacing two glutamines resulted in the absence of Ca^2+^ conductance in the simulations but allow for a notable K^+^ conductance (summarised in Fig. 8). When four arginines were present, Ca^2+^ conductance is still blocked as expected and K^+^ conductance is in the order of femtosiemens. These computational results are in remarkable agreement with patch clamp data on heteromeric channels incorporating edited GluA2 and have a much reduced unitary conductance that is difficult to resolve with conventional single-channel techniques (7). Of note, GluA2(R) homomers do conduct monovalent ions when in complex with the auxiliary subunit Stargazin (Stg) (22), which we did not test here. The degree to which currents from GluA2 (R) with Stg are due to rescue of plasma membrane trafficking by Stg is unclear, and perhaps unitary conductance is not altered much. Modification of pore properties by auxiliary subunits will be an important subject for future work (30). Our simulation results further revealed that the Ca^2+^ conductance is more sensitive to the number of arginines at the Q/R site than that of K^+^. On a naive level, our computational finding confirmed that ion conductance and selectivity depend strongly on the charge density in the narrow restriction site in the pore. However, the atomistic basis for this dependence, being that potassium ions can take a secondary route to pass the on-axis arginine side chains is most unexpected (movies S2 and S3, see Fig. 8). Overall, the multiple sites and multiple routes go some way to explaining the deviations of AMPARs from the Goldman-Hodgkin-Katz formulation of independence in physiological conditions(45). In our simulations, we used an open-state homomeric GluA2 channel as a structural model, gradually increasing the arginine level at Q/R site in different setups. Despite the general good agreement between the experimental and computational observations on conductance and cation selectivity of the Q- and R-forms of GluA2, it should be noted that introducing asymmetric arginine editing into homomeric GluA2 may not be feasible experiments. Therefore, comparison between the asymmetric R-edited GluA2 channel we have used here and heteromeric channels containing GluA2 and other GluA subunits should be made with caution.

**Fig. 8.**
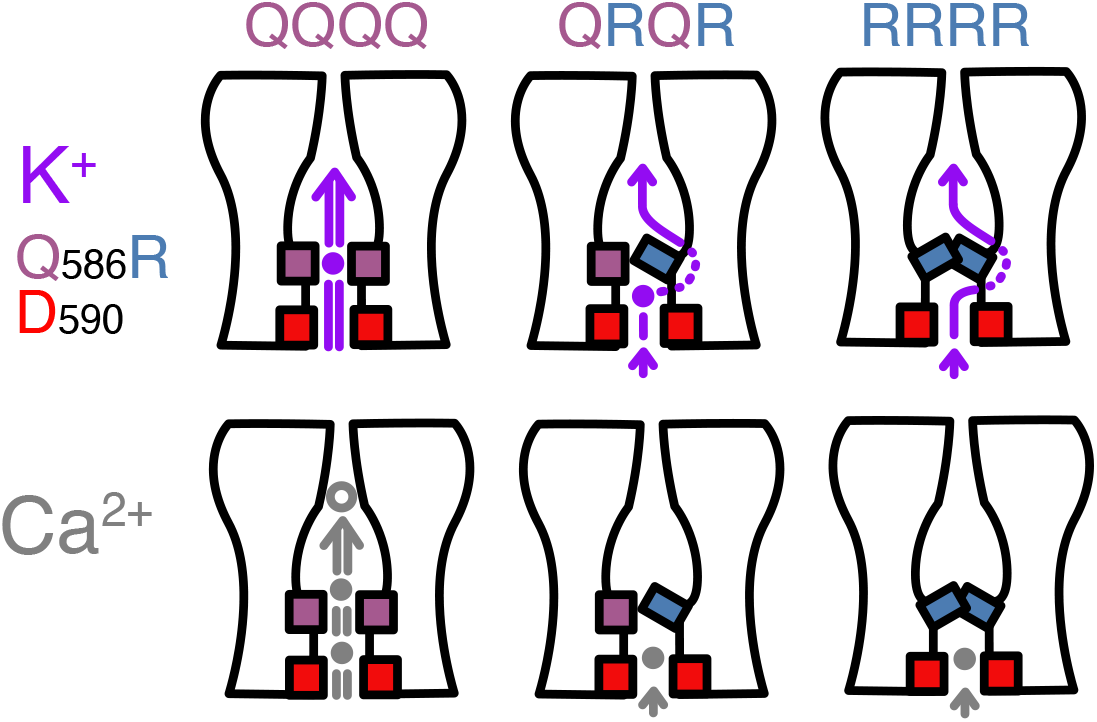
Cartoon representation of K^+^ and Ca^2+^ outward permeation through AMPARs with different levels of Q/R editing. Two of the four subunits are drawn. Q and R residues are shown as a mauve square and blue rectangle, respectively, the D590 residue as a red square. Filled circles represent binding sites, the open gray circle represents *site-G*, a weakly occupied Ca^2+^ binding site seen in Cryo-EM. Arrows and dotted paths represent the ion permeation routes. The double arrows represent rapid permeation.

We further observed in the simulations of mixed potassium and calcium a slowdown of K^+^ transit through the pore, caused by strong binding of Ca^2+^ at the intracellular side in the CDI region, leading to an inefficient knock-on mechanism between monovalent and divalent cations (movie S5). This effect was not observed between two monovalent cation types, which co-permeated through the channel pore with similar rates (movie S4). This finding replicates the previous experimental observations of Ca^2+^ blocking monovalent cation conduction (6). Notably, in previous MD simulations of ryanodine receptors with mixed K^+^ and Ca^2+^ ion types, the small-sized and doubly-charged Ca^2+^ were preferred in the confined region of the SF, while the energy barrier of K^+^ permeation increased in the presence of Ca^2+^ (46). Based on those simulations, it was suggested that charge and space competition is the major mechanism for high calcium selectivity in the Ryanodine receptor. Although we observed a similar mechanism in our simulations at relatively high Ca^2+^ concentrations, at low Ca^2+^/K^+^ ratio, both Ca^2+^ and K^+^ could occupy the filter with similar populations. Therefore, at physiological ion concentration where K^+^ concentration largely exceeds Ca^2+^ concentration, we propose that charge and space competition is not the major mechanism for the slowdown of monovalent cation conduction. Rather, Ca^2+^ binding at the intracellular CDI site blocks monovalent cation conduction.

Intriguingly, parallel functional comparison of NMDA, AMPA and Kainate receptor (KAR) Ca^2+^ to Na^+^ permeability gives strikingly different results, even between calcium-permeable AMPAR and KAR (45). These differences are despite the similar pore lining sequences. Open channel structures are now available for both NMDAR (47) and KAR (48) which should enable a computational investigation of this question across the entire family of iGluRs.

## Data availability

The simulation run input files, comprising the starting configuration and all necessary parameters for performing the MD simulations are deposited in Zenodo under https://doi.org/10.5281/zenodo.13918513

## Materials and Methods

### Patch clamp electrophysiology

The D590S and D590N mutants were made by overlap PCR in the BiQG.Ex4 I-E pRK5 vector. This vector has a CMV promoter followed by rat GluA2 (flip variant, unedited at the QR site, but edited at the RG site) and includes eGFP after an internal ribosomal entry site (IRES). Both constructs were confirmed by double-stranded DNA sequencing. The GluA2 WT subunit and mutants were introduced to HEK-293 cells (3 μg plasmid DNA per 35 mm dish) by transient transfection (calcium phosphate method (49)) and patch clamp experiments were performed 24-72 h later.

Two different extracellular solutions were used. The first was a standard NaCl buffer, containing (in mM): 150 NaCl, 0.1 MgCl2, 0.1 CaCl2, 5 HEPES. The pH was titrated to 7.3 with NaOH. The second buffer contained 100 mM CaCl_2_, titrated by CaOH also to pH 7.3. Patch pipettes with a tip resistance of around 3-5 MΩ were filled with intracellular solution containing (in mM): 115 NaCl, 10 NaF, 5 Na_4_BAPTA, 10 Na_2_ATP, 1 MgCl_2_, 0.5 CaCl_2_, 5 HEPES, titrated to pH 7.3 with NaOH. The internal solution was filtered with a 0.2 *μ*m Nylon filter and stored at –20º C in 0.5 mL aliquots. Once thawed it was kept on ice. Fluorescent cells were visualized on an IX-71 microscope equipped with a 470 nm LED. Outside-out patches were lifted into the outflow of a custom four barrel perfusion tool (raw glass from Vitrocom, V/6699, see (50) for details.) Solutions containing glutamate (10 mM) were applied by stepping a piezo translator (Physik Instrument) back and forth to move the interface between flowing control and glutamate solutions. Note that glutamate solutions (both Na^+^-rich and Ca^2+^-rich) used a glutamate stock titrated with NaOH and therefore contained at least 7.5 mM Na in excess.

Patch currents were recorded using a Axopatch 200B amplifier and digitised with an ITC-18 interface under the control of Axograph software. Responses were analysed using python code. The analysis code is publicly available at https://github.com/AGPlested/GluA2-D590X-Mutants. Data were plotted with Matplotlib. Patches were subjected to brief jumps into glutamate during a voltage ladder protocol with 10 mV steps from -90 mV to 90 mV. This command voltage was applied to each patch in both Na^+^-rich and Ca^2+^-rich conditions. Only patches with paired recordings in both conditions were included in the analysis. All traces were baseline corrected and Gaussian filtered at 1 kHz (using the ASCAM package, https://github.com/AGPlested/ASCAM).

Considering the biionic condition, and assuming that Na^+^ outside and Ca^2+^ inside were close to zero, the permeability ratio is given by the following equation:

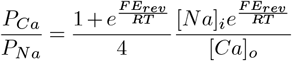

where F,R,T have their usual meanings and E_rev_ is the change in reversal potential from the pure sodium condition (minimal sodium gradient) This equation is derived from the GHK formalism. Note that, for Ca^2+^ permeation of AMPARs, deviations from the GHK assumptions were previously reported (45).

### MD simulations

The starting structure for the MD simulations was taken from an open-state cryo-EM structure of the GluA2 homo-tetramer (PDB: 5WEO (14)). The stargazin molecules were deleted, and the extracellular domains (ligand-binding and amino-terminal domains) were manually removed using PyMol (51). The truncation sites were at I504, G774, I633, and A820. We capped the C-termini with NME and the N-termini with ACE. The amino acid exchange from Q to R at the Q/R sites for the respective setups was also manually performed with PyMol. The program MODELLER (52) was used to model the missing intra-cellular loops, which were not resolved experimentally. For our all-atom MD simulations, we employed the GROMACS software suite (53) version 2019.6. The protein was parameterized with the C_HARMM_36 force field. The simulation setup was constructed with the C_HARMM_-GUI Membrane Builder (54–56): We embedded the protein in a POPC lipid bilayer, added ions (200 mM KCl, 100 mM CaCl2 in single cationic conditions, see Table S1), and solvated the system in TIP3P water. We used the standard C_HARMM_36 force field parameters for K^+^, Na^+^, and Cl^*−*^, while the multisite Ca^2+^ model (57) was ultilized in the simulations of calcium permeation. Energy minimization and equilibration were performed with the default C_HARMM_-GUI script provided with the simulation input files. For production runs we used the Computational Electrophysiology protocol (58) implemented in GROMACS. We generated a double bilayer setup based on the equilibrated single-bilayer system and equilibrated the new system for another 20 ns. We introduced a charge imbalance of 6 e, which was maintained throughout the simulations. This resulted in a transmembrane voltage between 300 and 600 mV (Figure S3). The protein and the membrane were stable in all our simulations (RMSD replot, Figure S4). Since we truncated the extracellular domains of the receptor, we applied position restraints on the last heavy atoms (non-H) of the peptide chain termini to prevent the ion channel from closing. The temperature was held at 303 K using the v-rescale thermostat. We used a Berendsen barostat with surface tension set at 250 bar nm (C_HARMM_-GUI: 25 dyne/cm) in the membrane plane (x y) and 1 bar in the z direction. All bonds were constrained using the LINCS algorithm and the time step used in all simulations was 2 fs. Each run had a simulation time of 250 ns, and we simulated 5 or 10 replicates per setup (Table S1), depending on the expected ion permeation rate.

### Analysis and visualization of MD trajectories

GRO-MACS built-in tools were used for trajectory post-processing, potential, and RMSD calculations. VMD (59) and PyMol were used for visualization of the MD trajectories. For the quantitative analysis of the trajectories, we wrote custom Python code, using the MDAnalysis package (60, 61) for analysis and the matplotlib library (62) for data visualization. For the analysis of ion conduction, one ion permeation event was recorded when the ion passed through the entire channel pore.

## ACKNOWLEDGEMENTS

We thank Miriam Lohr for preparing the plasmids. This work was funded by the Leibniz-Forschungsinstitut für Molekulare Pharmakologie (FMP) and the Deutsche Forschungsgemeinschaft through the Research Unit 2518 DynIon, project P03, to A.J.R.P. and H.S and under Germany’s Excellence Strategy (EXC-2049-390688087-NeuroCure) to A.J.R.P.. The authors gratefully acknowledge the computing time made available to them on the Erlangen National High Performance Computing Center (NHR@FAU) and high-performance computer “Lise” at the NHR Center NHR@ZIB.

## Supporting Information

**Movie S1.** A representative trajectory showing Ca^2+^ permeation (grey spheres) through the GluA2(Q) channel in MD simulation (trajectory is from (28)), overlaid with experimentally-determined Ca^2+^ binding sites from a Cryo-EM study (orange mesh) (24)

**Movie S2.** Part of a representative trajectory of the RRRR setup with potassium ions. An ion enters an alternative, “off-axis” side pathway, and will eventually permeate the entire channel.

**Movie S3.** Later section of the same representative trajectory of the RRRR setup as in Movie S2. The permeating potassium ion eventually leaves the side pathway and permeates the channel through the gate.

**Movie S4.** A representative trajectory of mixed monovalent cation permeation through the GluA2(Q) channel. K^+^(magenta) and Na^+^(blue) were present at a ratio of 1:1

**Movie S5.** A representative trajectory of mixed cation permeation through the GluA2(Q) channel. K^+^(magenta) and Ca^2+^(grey) were present at a ratio of 5.8:1.

**Table 1.**
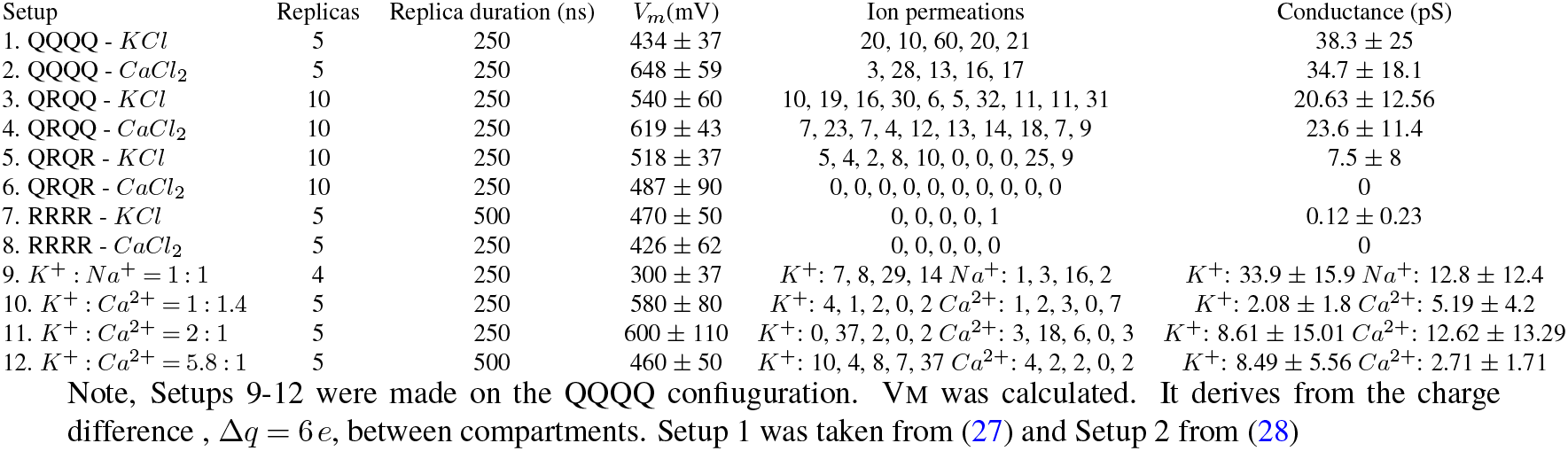
Summary of computational electrophysiology simulations.

**Table 2.** Details of the simulation setups.

| Setup | Total $N$ of atoms | Total $N$ of $\text{H}_2\text{O}$ | Cation concentration ($\frac{\text{mmol}}{\text{L}}$) | total $N$ of POPC | Box dimensions ($\text{\AA}$) |
| --- | --- | --- | --- | --- | --- |
| 1. QQQQ - $\text{KCl}$ | 229412 | 49184 | 213.6 | 444 | 97.6 x 97.6 x 230.7 |
| 2. QQQQ - $\text{CaCl}_2$ | 187698 | 35234 | 106.7 | 444 | 96.7 x 96.7 x 197.0 |
| 3. QRQQ - $\text{KCl}$ | 209924 | 43696 | 179.7 | 436 | 96.8 x 96.8 x 216.6 |
| 4. QRQQ - $\text{CaCl}_2$ | 210270 | 43696 | 88.6 | 436 | 96.5 x 96.5 x 217.8 |
| 5. QRQR - $\text{KCl}$ | 210054 | 43734 | 179.6 | 436 | 95.8 x 95.8 x 221.6 |
| 6. QRQR - $\text{CaCl}_2$ | 211434 | 43920 | 88.1 | 438 | 95.3 x 95.3 x 224.6 |
| 7. RRRR - $\text{KCl}$ | 210916 | 43920 | 181.3 | 438 | 96.1 x 96.1 x 221.1 |
| 8. RRRR - $\text{CaCl}_2$ | 211264 | 43961 | 90.7 | 438 | 95.6 x 95.6 x 223.3 |
| 9. $\text{K}^+ : \text{Na}^+ = 1 : 1$ | 229412 | 49184 | $\text{K}^+$: 106.8, $\text{Na}^+$: 106.8 | 444 | 97.6 x 97.6 x 230.7 |
| 10. $\text{K}^+ : \text{Ca}^{2+} = 0.7 : 1$ | 229762 | 49184 | $\text{K}^+$: 56.2, $\text{Ca}^{2+}$: 78.7 | 444 | 96.2 x 96.2 x 239.7 |
| 11. $\text{K}^+ : \text{Ca}^{2+} = 2 : 1$ | 229652 | 49184 | $\text{K}^+$: 105.7, $\text{Ca}^{2+}$: 54.0 | 444 | 95.9 x 95.9 x 241.3 |
| 12. $\text{K}^+ : \text{Ca}^{2+} = 5.8 : 1$ | 213738 | 44050 | $\text{K}^+$: 145.6, $\text{Ca}^{2+}$: 25.1 | 440 | 97.0 x 97.0 x 219.7 |

Note, Setup 1 was taken from (27) and setup 2 from (28). Setups 9-12 were made on the QQQQ configuration.

**Fig. 9.**
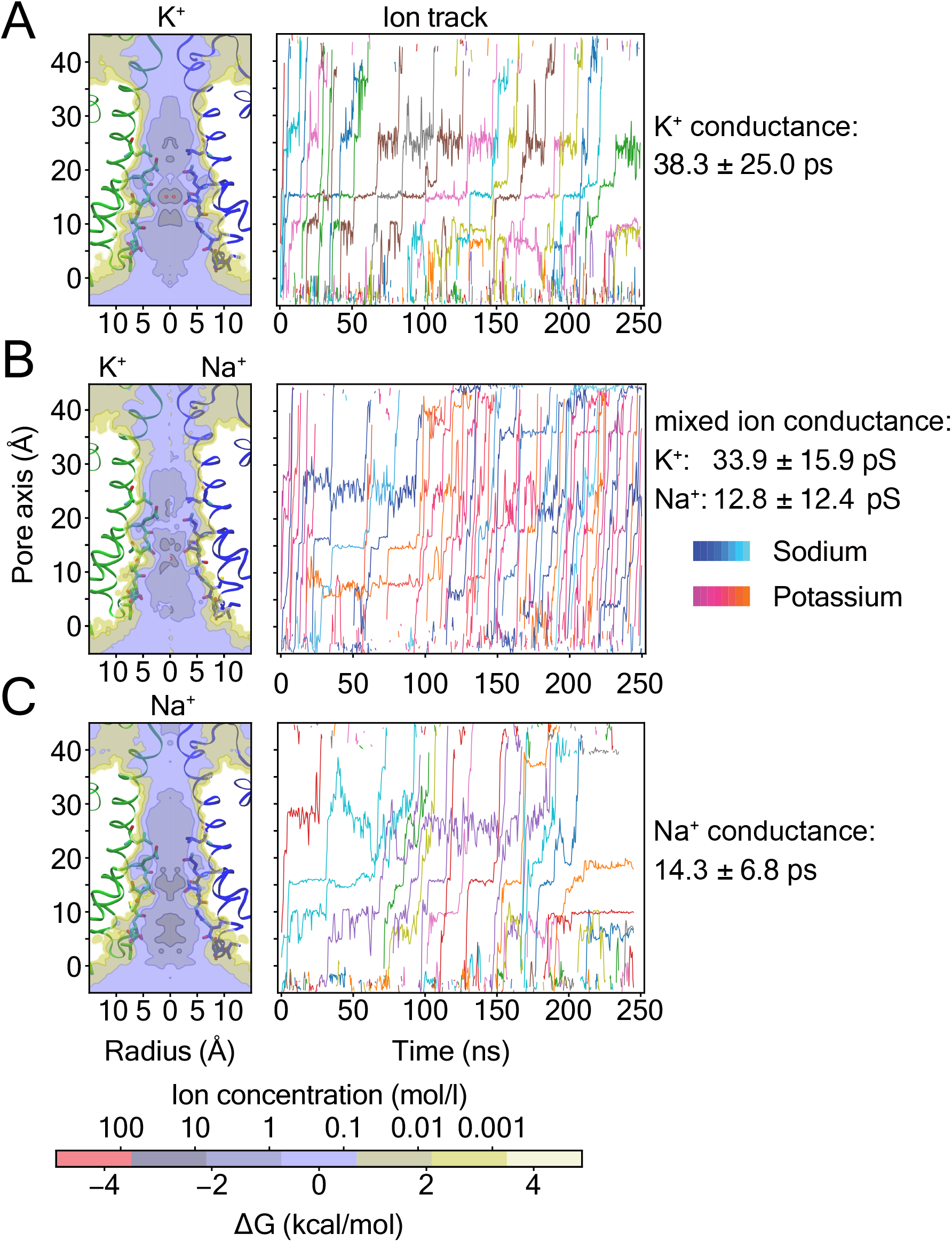
Comparison of two-dimensional ion occupancy (left) and representative ion track plots (right) for simulations of (A) K^+^ only, (B) a mixed system of K^+^ (reddish lines) and Na^+^ (bluish lines), and (C) Na^+^ only. The data for parts A and C are from (27).

**Fig. 10.**
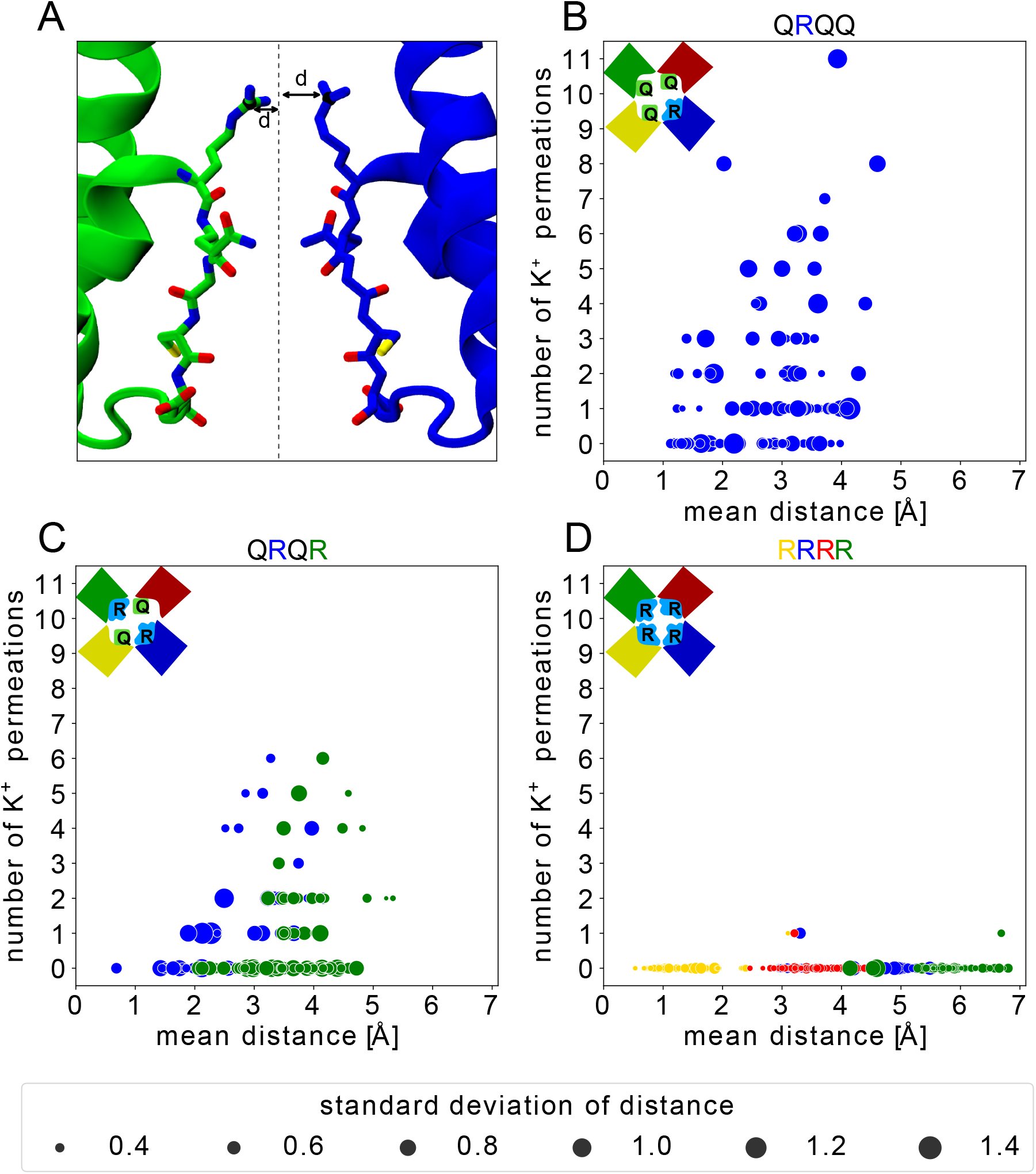
(A) Minimum distances of R586 from the central channel pore aixs (*d*). Correlation between instantaneous ion permeation and the average d within each 25 ns simulation interval for the (B) QRQQ, (C) QRQR, and (D) RRRR setups.

**Fig. 11.**
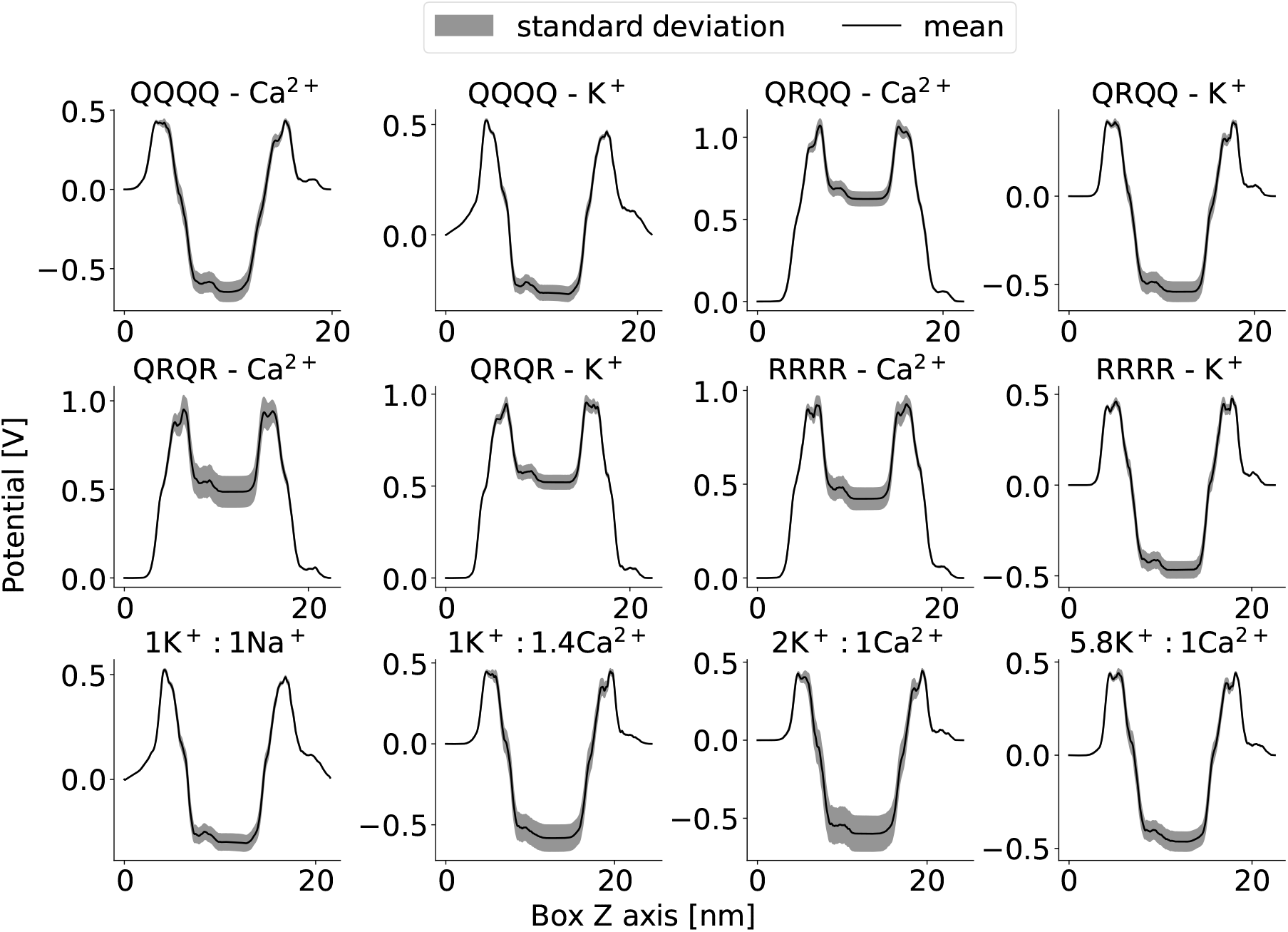
Averaged potential profiles along the z-axis (channel axis) for all simulation setups.

**Fig. 12.**
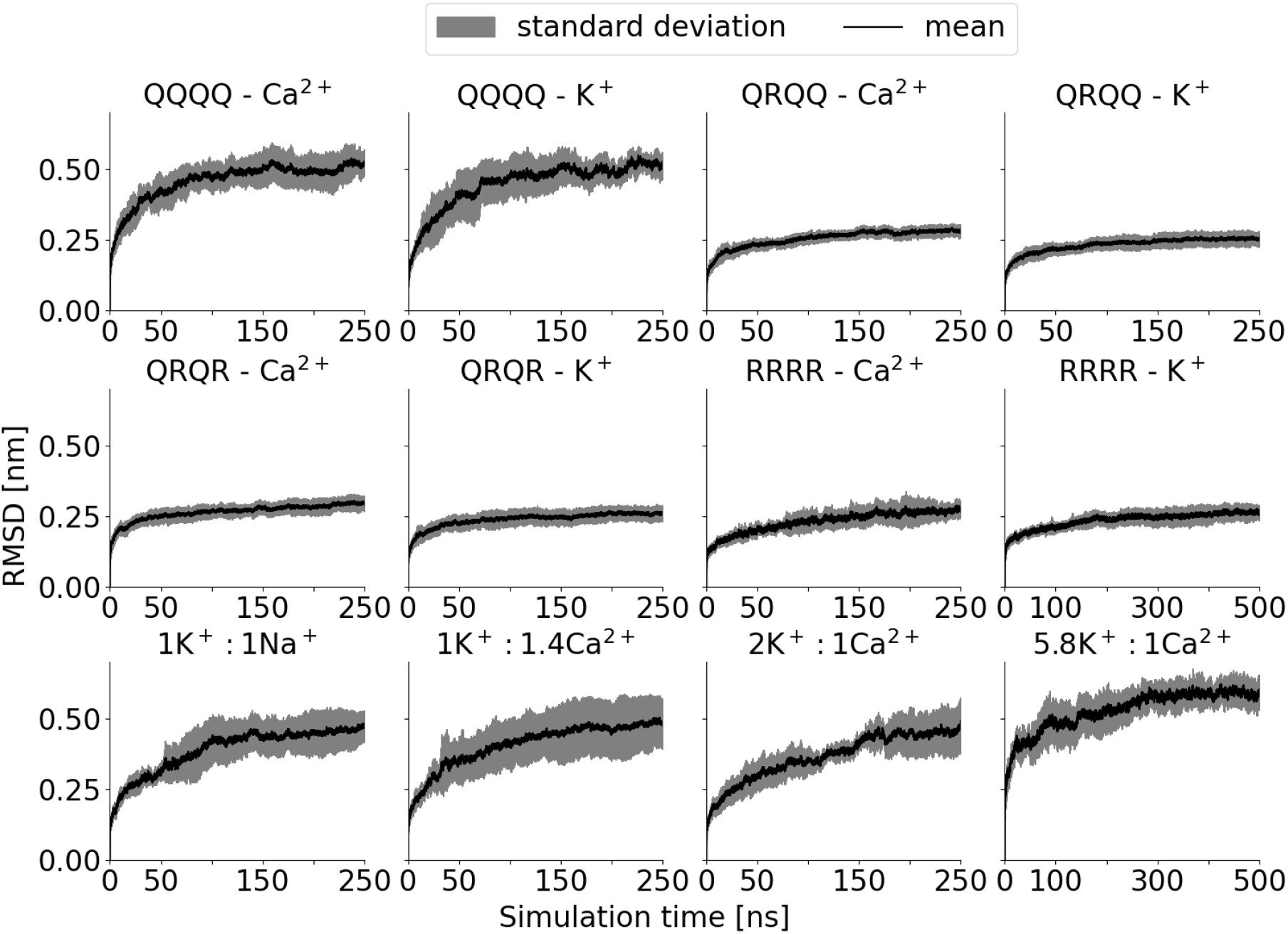
The average RMSD values (black lines) and standard deviations (shaded areas) of the C*α* atoms of the AMPA receptor pore domain, across all simulation setups.

